# Identification of protein secretion systems in bacterial genomes

**DOI:** 10.1101/031039

**Authors:** Sophie S Abby, Jean Cury, Julien Guglielmini, Bertrand Néron, Marie Touchon, Eduardo PC Rocha

**Author notes:** Division of Archaea Biology and Ecogenomics, Department of Ecogenomics and Systems Biology, University of Vienna, A-1090 Vienna, Austria (Sophie S Abby). Bioinformatics and Biostatistics HUB, Center of Bioinformatics, Biostatistics and Integrative Biology (C3BI), Institut Pasteur, Paris, 75015, France.

## Abstract

Bacteria with two cell membranes (diderms) have evolved complex systems for protein secretion. These systems were extensively studied in some model bacteria, but the characterisation of their diversity has lagged behind due to lack of standard annotation tools. We built models for accurate identification of protein secretion systems and related appendages in bacteria with LPS-containing outer membranes. They can be used with MacSyFinder (standalone program) or online (http://mobyle.pasteur.fr/cgi-bin/portal.py#forms::txsscan). They include protein profiles and information on the system’s composition and genetic organisation. They can be used to search for T1SS-T6SS, T9SS, and accessorily for flagella, Type IV and Tad pili. We identified ~10,000 systems in bacterial genomes, where T1SS and T5SS were by far the most abundant and widespread. The recently described T6SS^iii^ and T9SS were restricted to Bacteroidetes, and T6SS^ii^ to *Francisella.* T2SS, T3SS, and T4SS were frequently encoded in single-copy in one locus, whereas most T1SS were encoded in two loci. The secretion systems of diderm Firmicutes were similar to those found in other diderms. Novel systems may remain to be discovered, since some clades of environmental bacteria lacked all known protein secretion systems. Our models can be fully customized, which should facilitate the identification of novel systems.Introduction

## Introduction

Proteins secreted by bacteria are involved in many important tasks such as detoxification, antibiotic resistance, and scavenging ^1^. Secreted proteins also have key roles in both intra- and inter-specific antagonistic and mutualistic interactions ^2,3^. For example, they account for many of the virulence factors of pathogens ^4,5^. Bacteria with a LPS-containing outer-membrane (abbreviated “diderm-LPS” in this article) require specific protein secretion systems. Six types of secretion systems, numbered T1SS to T6SS, were well characterised by numerous experimental studies (for some general reviews see ^6-8^). The T9SS (PorSS) was more recently uncovered in Bacteroidetes ^9,10^. In this study, we focus only on typical diderm-LPS protein secretion and homologous systems. A few other systems have been described in the literature. For example, the chaperone-usher pathway, sometimes named T7SS, and the T8SS were not included in this study because they are only involved, respectively, in the export of type I pili and curli ^11^. The ESX system of *Mycobacteria,* T7SS according to some authors ^12^, was also ignored because it is absent from diderm-LPS bacteria.

The role of secreted proteins has spurred interest in the production of ontologies to categorise ^13^, and computational methods to identify and characterise protein secretion systems (Table 1). This is a difficult task for a number of reasons. Firstly, protein secretion systems are large machineries with many different components, some of which are accessory and some interchangeable. Secondly, many of their key components are homologous between systems, which complicates their discrimination. For example, T2SS, T4SS and T6SS include distinct but homologous NTPases ^14^. Some bacterial appendages require their own secretion systems to translocate their extracellular components ^15,16^, and these are sometimes partly homologous to classical secretion systems. For example, several components of the type IV pilus (T4P) and the Tad pilus are homologous to components of the T2SS from *Klebsiella oxytoca* ^17^. Thirdly, the sequences of secreted proteins, including extracellular components of the secretion systems, evolve rapidly, thereby complicating the identification of homology by sequence similarity ^18^. Fourthly, loci encoding secretion systems are frequently horizontally transferred and lost ^19,20^, leading to the presence of partial (often inactive) systems in genomes ^21^. Finally, experimental studies have focused on a small number of occurrences of each type of system, complicating the assessment of their genetic diversity. On the other hand, secretion systems are often encoded in one or a few neighbouring operons made of highly conserved, and in some cases highly specific, components. This information can facilitate the identification of genes encoding secretion systems in genome data ^22^,^23^.

**Table 1.**
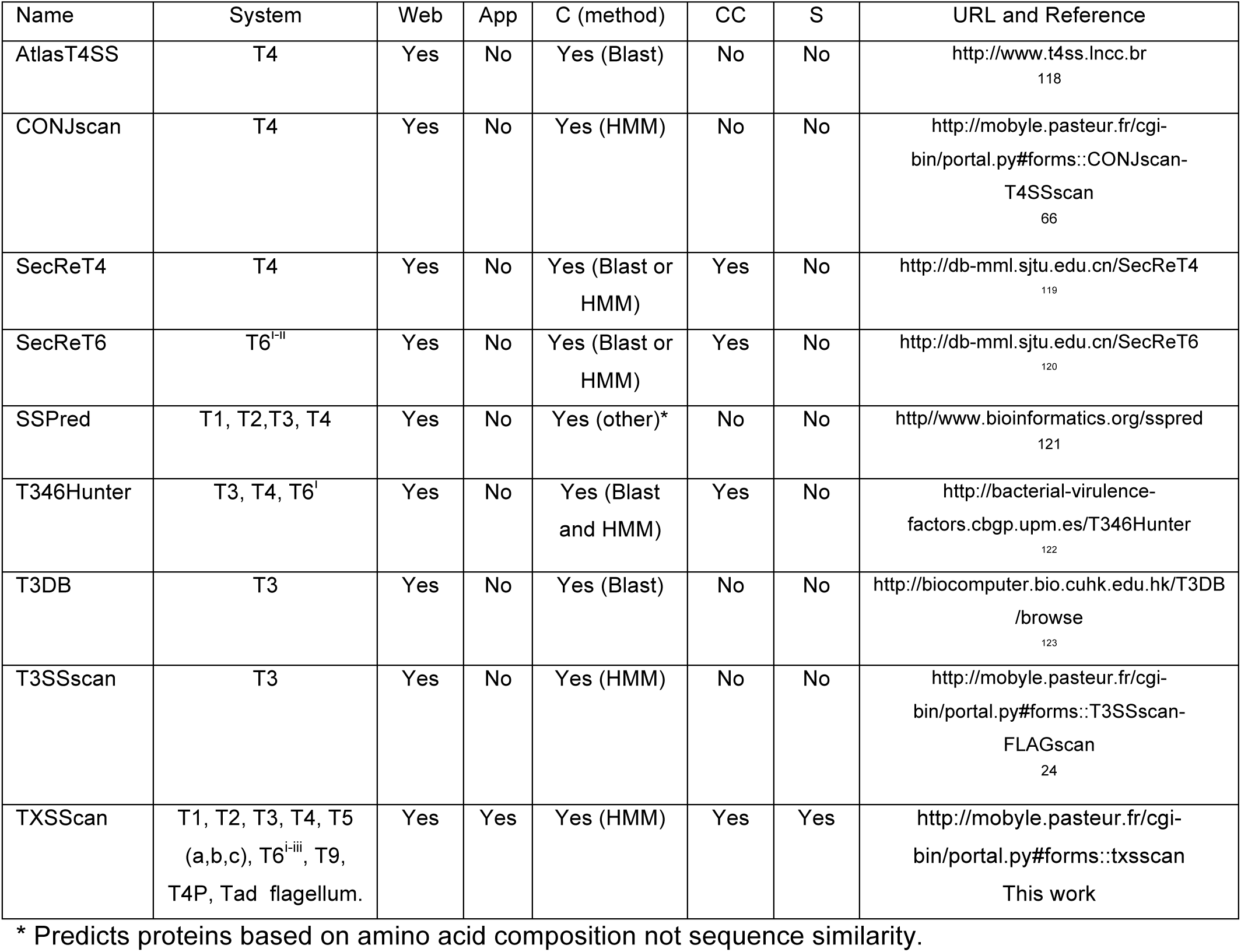
Public webservers (Web) and downloadable applications (App) to identify components (C), clusters of components (CC), or complete (eventually scattered, S) bacterial protein secretion systems.

Several programs were previously made available to identify components of some, but not all, protein secretion systems (Table 1). These programs are very useful to the biologist interested in browsing the known systems or in annotating a small set of sequences. However, they are web-based, and thus poorly adapted for the analysis of very large datasets. Most of them were not designed to categorise systems as complete or incomplete, and the few that were do not allow the identification of systems scattered in the chromosome, nor the re-definition of the categorisation parameters. Yet, the ability to re-define the type and number of components in a system facilitates the search for distant variants of the well-known experimental model systems ^24^. We have used the vast body of knowledge accumulated from experimental studies of model protein secretion systems to identify new occurrences of these systems. This was achieved by building computational models that account for the systems’ composition and genetic organization. The models can be plugged in MacSyFinder ^25^ to identify protein secretion systems. This can be done using the standalone application and using local resources, or using the webserver version available at http://mobyle.pasteur.fr/cgi-bin/portal.py#forms::txsscan. The results can be visualized with MacSyView ^25^. In the standalone application, the users can easily modify the models to change the composition and genetic organisation of the secretion systems. Some of these parameters can also be modified in the webserver version. The accuracy of the models was quantified against an independent dataset of experimentally validated systems. Importantly, we provide models to search for an unparalleled number of protein secretion systems (and some partly homologous systems): T1SS, T2SS (Tad and T4P), T3SS (flagellum), T4SS (conjugation system), T5SS, T6SS^i-iii^, and T9SS. We used the models to search for protein secretion systems in a large panel of bacterial genomes. Previous surveys, mostly dating from a time when few genomes were available, analysed the distribution of some specific protein secretion systems in genomes or metagenomes ^17,20,24,26-31^. Due to space limitations, we will not attempt at re-assessing all these works. Instead, we describe our models, show their accuracy, and use them to provide a broad view of the relative distribution of the different protein secretion systems.

## Results and discussion

### Overview of the approach

We defined 22 customisable models for the protein secretion systems and related appendages (File S1, Fig. 1-6, Fig. S1-S5). This was done in four steps (see Materials and Methods for details). Firstly, we searched the primary literature, reviews and books for references of well-studied systems ^9,16,20,26,28,32-40^. We used them to define two independent datasets (*reference* and *validation*) of experimentally studied secretion systems (Tables S1 and S2). Secondly, the *reference dataset* was used to define the model for each type of system. The model includes information on the number of components that are *mandatory* (necessarily present in a system), *accessory* (not necessarily present in the system), and *forbidden* (never present in the system). Each component is associated with a protein profile that is used to search its occurrences (with HMMER ^41^). Our models use 204 protein profiles (included in the package), of which 194 were built in our laboratory and the rest taken from public databases (Table S4). Protein profiles are more sensitive and specific than Blast-based approaches ^42^. The model indicates which genes are co-localised (at less than a given distance relative to contiguous genes in the cluster), and which genes might be encoded elsewhere in the genome (designated “loners”). Thirdly, the models were validated both in the *reference* and in the independent *validation* datasets using MacSyFinder ^25^. Finally, the models were used with MacSyFinder to identify occurrences of each system in 1,528 complete genomes of diderm species. This procedure retrieved automatically all validly identified secretion systems. It also retrieved all hits to each component identified in the genomes whether they are part of a protein secretion system or not. In the following sections we describe the models of each type of protein secretion system and the occurrences of the system in bacterial genomes.

**Figure 1.**
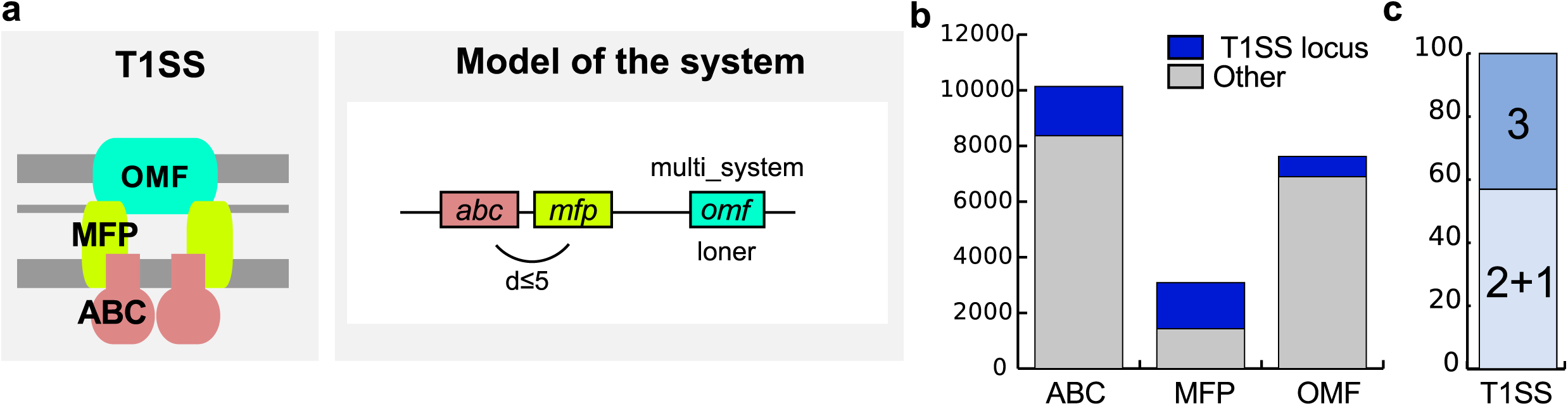
Model and results for the T1SS. **a.** Schema of the structure (left panel) and model of the genetic Organisation (right panel) of T1SS. We built protein profiles for the three components and modelled the two possible genetic architectures of the T1SS: one with the three components encoded in a single locus (*inter_gene_max_space* parameter in MacSyFinder: d<5 genes), another with the ABC transporter and the MFP encoded in a locus while the OMF is further away (*loner* attribute). A single OMF can also be used by different T1SS ^45^ and this is noted by the attribute *multi_system.* **b.** Distribution of hits for the protein profiles of the T1SS components, separated in two groups: hits effectively part of a T1SS main locus (*i.e.,* containing at least ABC and MFP, grey) and hits found elsewhere (“Other”, blue). Even if encoded outside of “main loci” (grey area of the bar), OMF might be involved in T1SS (*loner* property), whereas it is not the case for ABC and MFP. **c.** T1SS encoded in one single locus (ABC, MFP and OMF co-localise) (3) or in two (OMF encoded away from the other components) (2+1).

**Figure 2.**
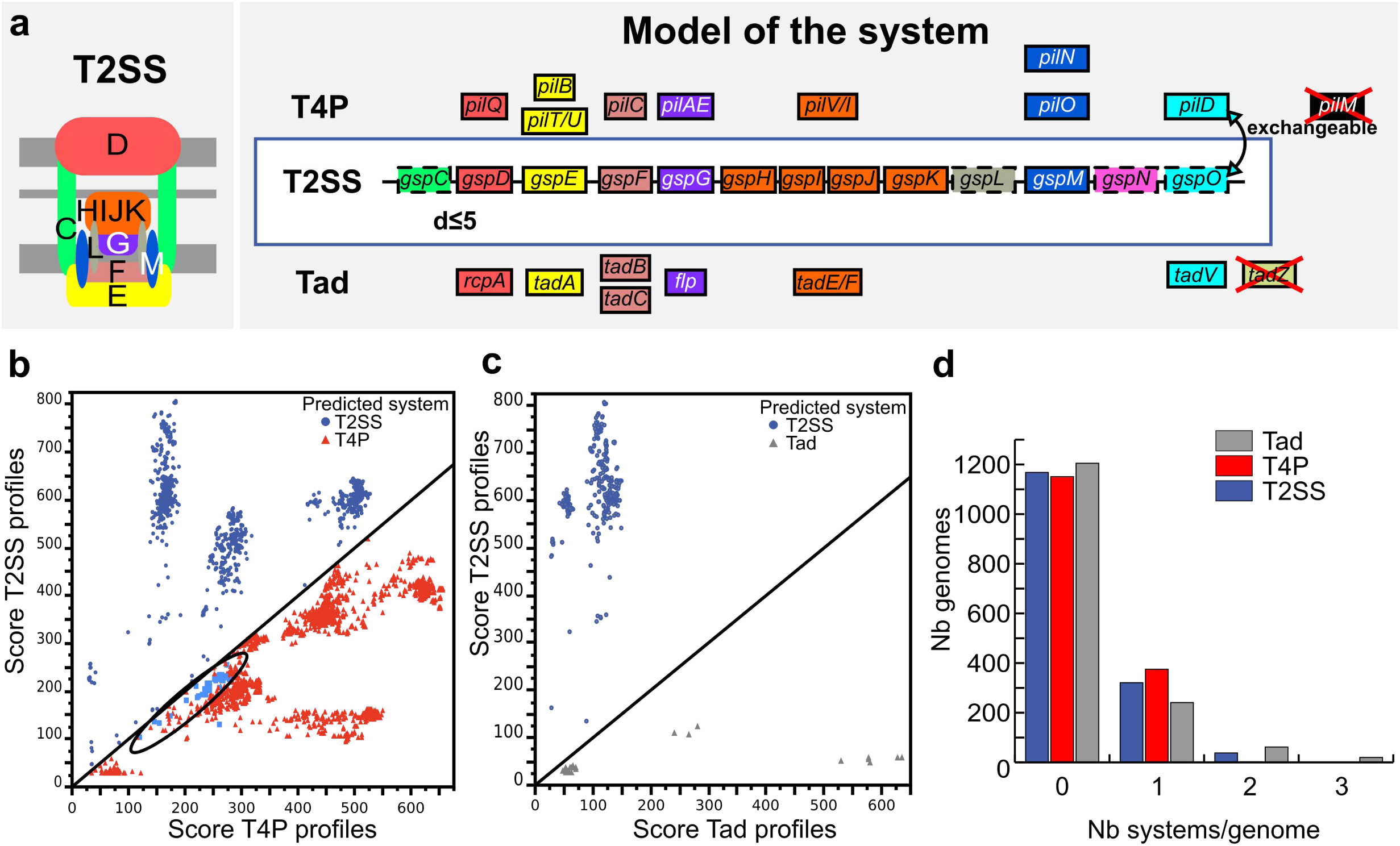
Model of the T2SS for detection and discrimination from the T4 and Tad pili. **a.** Schema of the structure (left panel) of T2SS, and model of its genetic organisation (right panel), indicating components with homologies with T4P and Tad pilus. We built protein profiles for all these components (Tables S4 and S5). Protein families represented by the same colour are homologous, and their profiles often match proteins from the other systems (except for the Flp and TadE/F families that are less similar). Some prepilin peptidases of T2SS and T4P are defined as functionally interchangeable ^108-110^ (curved double-headed arrow, exchangeable attribute). Boxes represent components: mandatory (plain), accessory (dashed) and forbidden (red crosses). **b.** Scores of proteins matched with the profiles of T2SS and T4P. The components of actual T2SS (dark blue) and actual T4P (in red) are well separated, indicating that in each case the best match corresponds to the profile of the correct model system. The exceptions (blue points surrounded by a black ellipse) concern the prepilin peptidases (light blue squares, circled in blue), which are effectively inter-changeable. **c.** Representation similar to **b**, but for the comparison between T2SS (blue) and Tad (grey) systems. In this case, the separation is perfect: the proteins always match better the protein profile of the correct system. **d.** Number of detected systems per genome among the 1,528 genomes of diderm bacteria.

**Figure 3.**
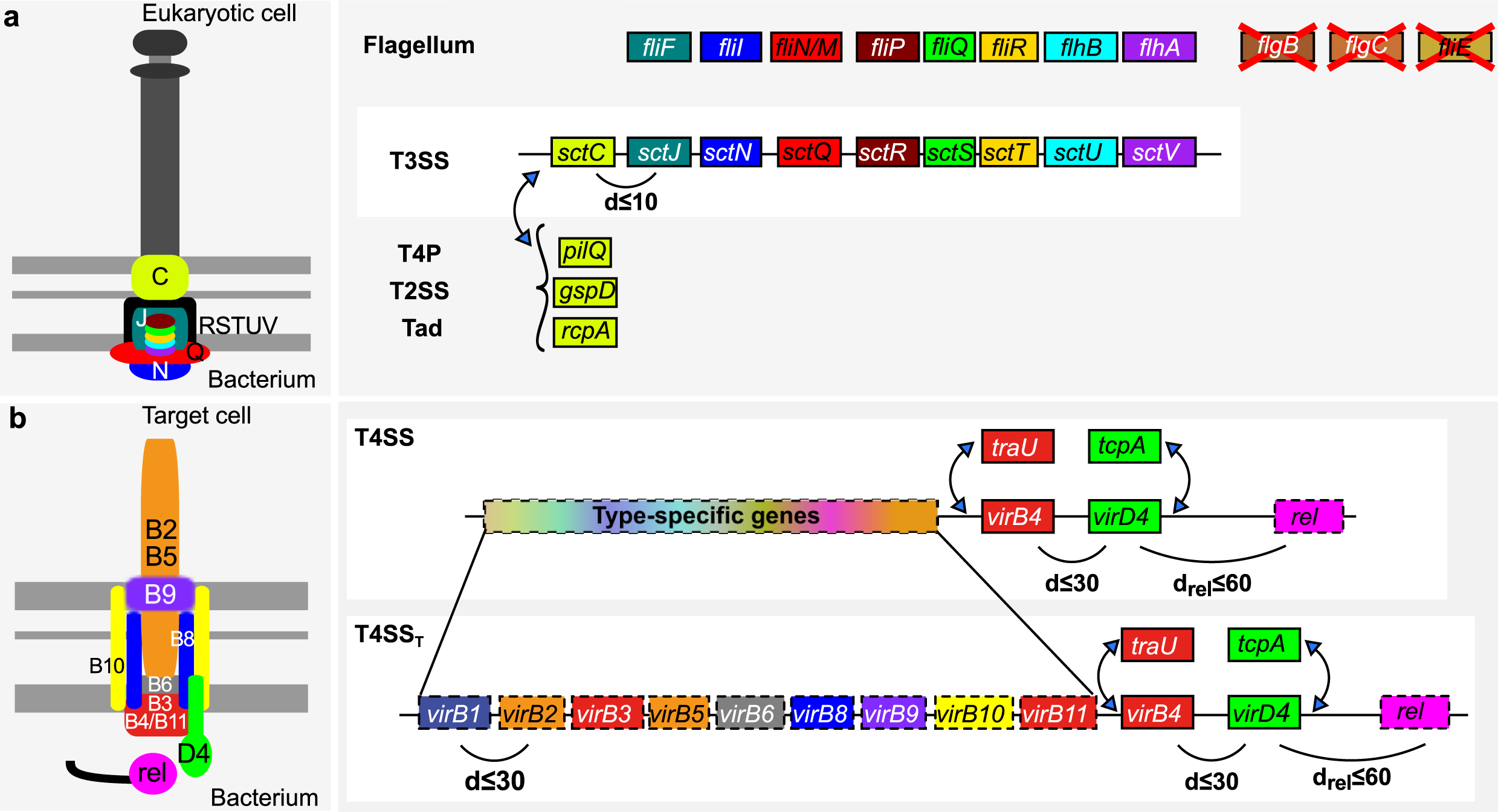
Model of T3SS and T4SS. **a.** The models of T3SS and flagellum were built based on a previous study ^24^ (representation conventions as in Fig. 2). Of the nine mandatory components for the T3SS only the secretin is forbidden in the model of the flagellum. Conversely, three flagellum-specific components are forbidden in the T3SS model. Three different types of secretins are found in T3SS derived from different appendages, which are thus defined as exchangeable in the model. **b.** Models of the T4SS were built based on a previous study Two different proteins have been described as type 4 coupling proteins (T4CP: VirD4 and TcpA) and two as the major ATPases (VirB4 and TraU, which are homologous). Some pT4SS lack a T4CP and secrete proteins from the periplasm ^64^ The relaxase (rel), is necessary for conjugation but not for protein secretion, although some relaxase-encoding T4SS are both cT4SS and pT4SS ^111-113^ Only two MPF types are associated with protein secretion - pT4SS_I_ and pT4SS_T_, corresponding to MPF_I_ and MPF_T_ types. The specificity of type-specific profiles is assessed in Fig. S2.

**Figure 4.**
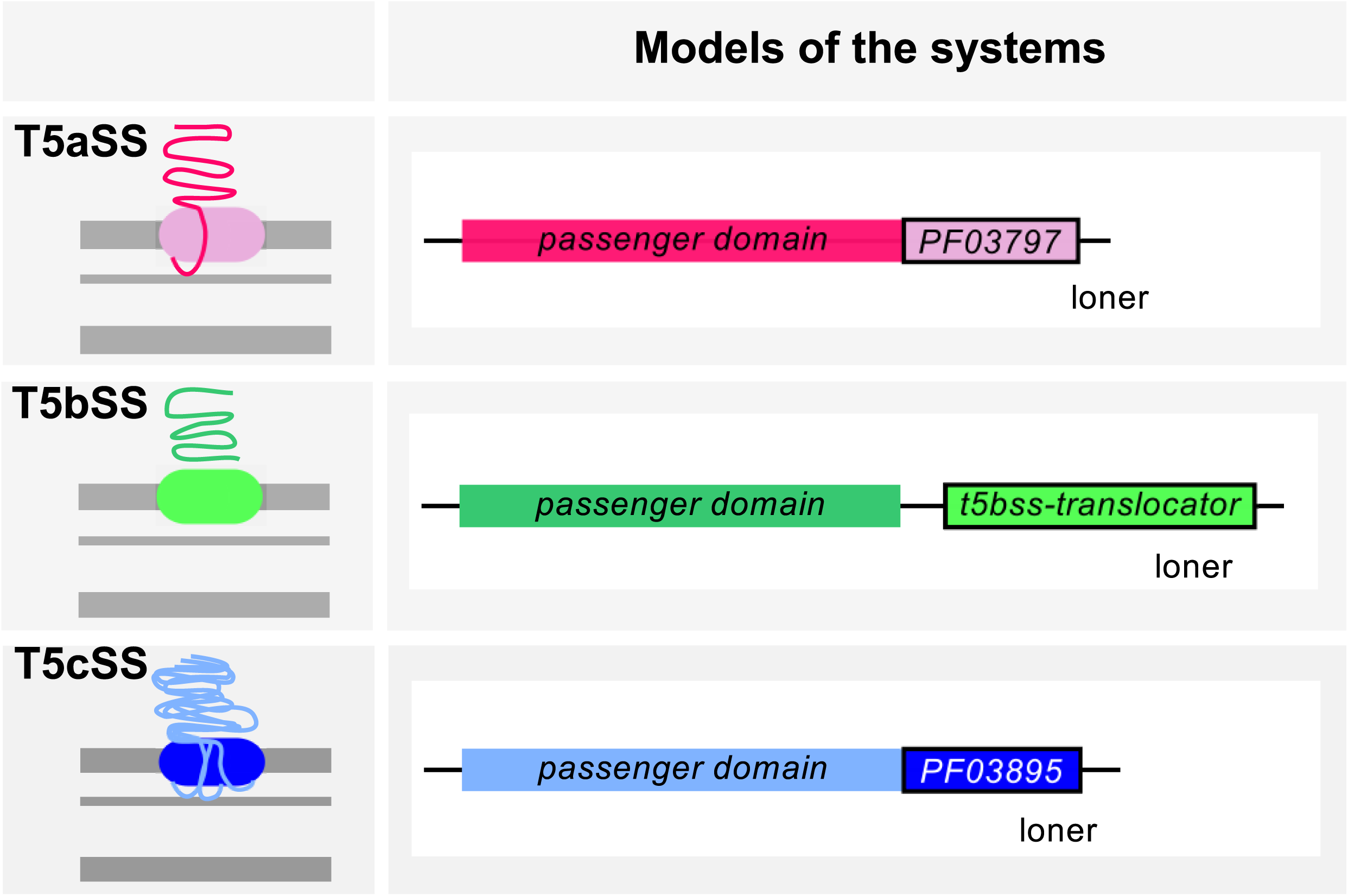
Model of the T5aSS, T5bSS and T5cSS. The left panel shows simplified schemas of the T5SS, and the right panel displays the respective genetic model (only one component that is classed as *loner*). The translocator, pore-forming domains were searched using PFAM domains for T5aSS and T5cSS (resp. PF03797 and PF03895), and a profile built for this work for the T5bSS (Tables S1, S4, S5).

**Figure 5.**
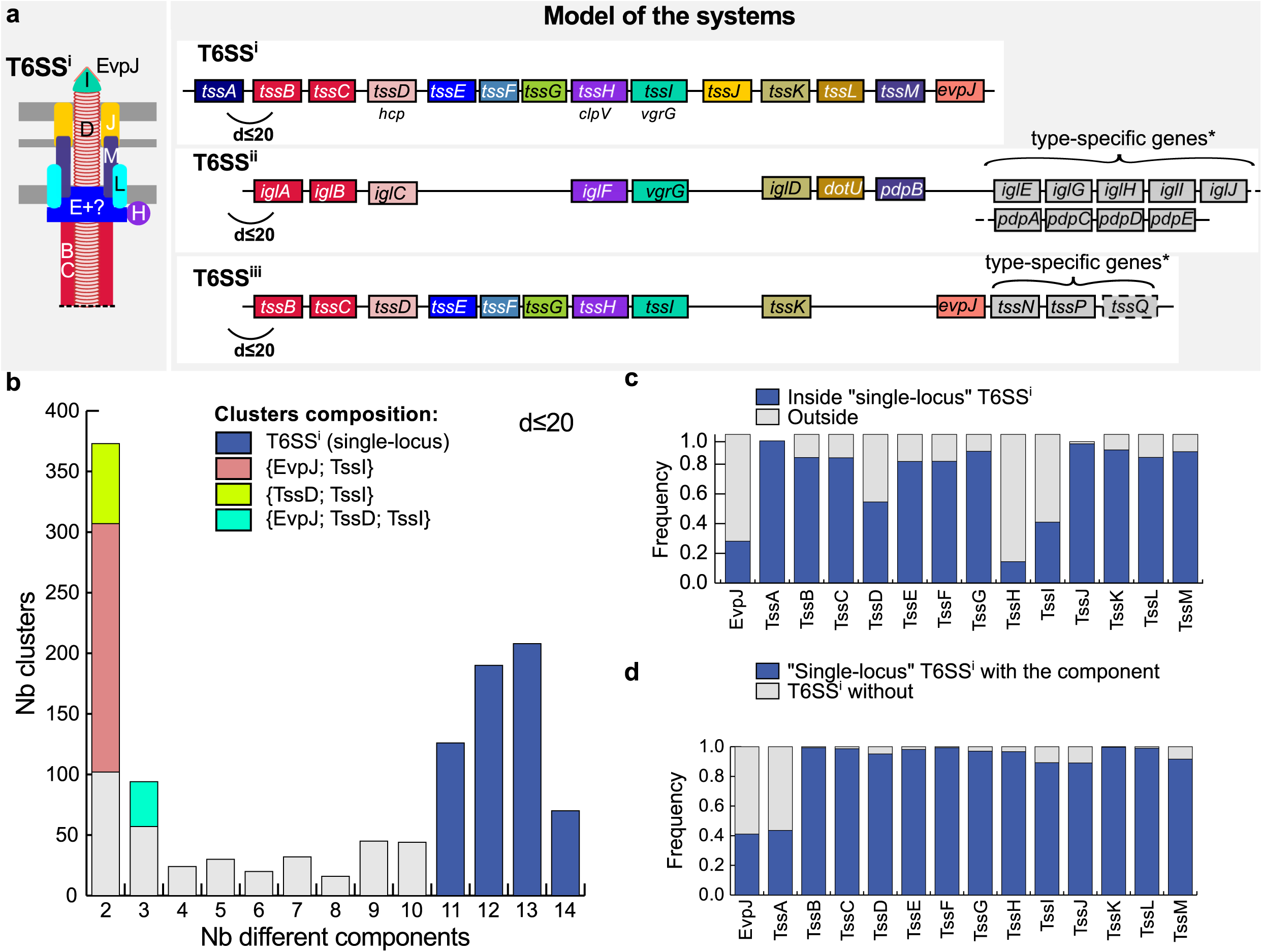
Model and results for the detection of T6SS. **a.** The left panel shows the schema of the structure of T6SS^i^, and the right panel displays the genetic model of the three sub-types of T6SS (representation conventions as in Fig. 2). For T6SS^i^, we built profiles for the 14 mandatory components, which were clustered if at a distance of d≤20 (see Fig. S4). For T6SS^ii^ and T6SS^iii^, we built 17 and 13 profiles respectively. All components were set as mandatory, except for TssQ, which is found in half of the T6SS^ii^. Homologies between components that are displayed by the mean of the same colours of boxes between the different sub-types are based on previous studies. *Putative type-specific genes are displayed in grey boxes that do not represent homologies. However, several putative homologies were retrieved using Hhsearch (e-value<1 and p-value<0.05) on T6SS^ii^ components: iglC (tssG), iglG (tssF), iglH (tssE), iglJ (tssH) and pdpD (tssH). **b.** Number of different components per cluster of T6SS^i^. Following this analysis, we set the quorum parameter of T6SS^i^ to 11. **c.** Frequency of hits for each type of T6SS^i^ components in the genomes. Hits matching a single-locus T6SS^i^ are in blue. The other hits match outside the T6SS^i^ loci. **d.** Frequency of each component within single-locus T6SS^i^. The components EvpJ and TssA were detected in less than 45% of the T6SS^i^, while the other components are found in most T6SS^i^ loci (>89%).

**Figure 6.**
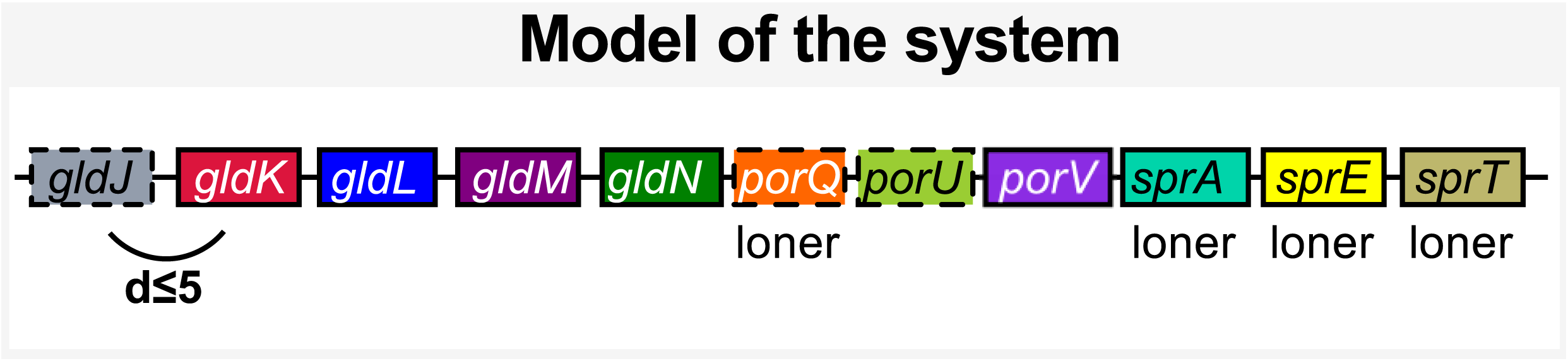
Genetic model of the T9SS. The representation follows the conventions of Fig. 2. The model includes 11 components for which 13 protein profiles were obtained from PFAM (SprA and SprA-2), TIGRFAM (GldJ, GldK, GldL, GldM, GldN and SprA-3) or designed for this study (PorU, PorV, PorQ, SprE, SprT). Four components were declared as *loners.* The co-localisation distance for the others was set at d≤5 (see Fig. S5). As several profiles were available for SprA, we included them all in the models, and declared them as exchangeable homologs in the model. GldJ is not part of the secretion system, but of the gliding motility system. It was included in the model as it facilitates the detection of T9SS that co-localises with it.

### T1SS

We built protein profiles for the three essential components of T1SS ^32,43,44^: the ABC-transporter (ATP-binding cassette transporter) providing an inner membrane channel, the porin (outer membrane factor, OMF) forming the outer-membrane channel, and the inner-membrane anchored adaptor protein (or membrane-fusion protein, MFP) that connects the OMF and the ABC components (Materials and Methods, Tables S4-S5). T1SS can be difficult to identify because its components have homologs involved in other machineries, *e.g.,* in ABC transporters for the ABC and in drug efflux systems for the MFP, or can themselves be involved in other machineries, in the case of the OMF ^45-49^. The design of the model was facilitated by the previous observation that genes encoding the ABC and MFP components are always co-localised in T1SS loci ^32,43^. The model is described in Figure 1a, and its use resulted in the correct identification of all T1SS in the reference and in the validation datasets. Overall, 20,847 proteins matched the protein profiles of the T1SS components in the bacterial genomes (Fig. 1b). The vast majority of these were not part of T1SS because they did not fit the T1SS model. We found 1,637 occurrences of the T1SS model in 821 genomes (Fig. 1b). The remaining proteins are probably associated with the numerous other systems carrying components homologous to those of the T1SS.

We found T1SS in more than half of the genomes of diderm bacteria (54%). Some genomes contained many systems; *e.g., Bradyrhizobium oligotrophicum* S58 and *Nostoc sp.* PCC 7524 encoded a record number of 9 systems (Table S3). ABC and MFP were encoded together and OMF apart in more than half (57%) of the T1SS. Hence, the co-localisation of the genes encoding the three components concerns a minority of T1SS (Fig. 1c). We found 95 loci encoding ABC and MFP in replicons lacking OMF. Many of these systems may be functional, since 94 of these loci were found in genomes encoding at least one OMF in another replicon. Such multi-replicon functional T1SS have been previously reported ^50^.

### T2SS, T4P and Tad pili

T2SS are encoded by 12 to 16 genes, many of which are homologous to components of the T4P and the Tad pilus ^17,51,52^ (Fig. 2). We used the protein families conserved in the reference dataset to build 13 protein profiles for T2SS, 11 for type IV pili and 10 for Tad pili (Materials and Methods, Tables S4-S5). No profiles were built for GspA and GspB, because they were rarely identified in T2SS plus we could not build protein families given low sequence similarity. The most frequent components in the reference dataset were defined as mandatory in the models, and the least conserved as accessory. Some profiles built for one type of system produced matches to (homologous) components of other types of systems. Discrimination between systems was facilitated by the definition of some specific components as forbidden (*e.g.,* GspC was declared forbidden in Tad and T4P).

Using the models, we correctly identified all of the T2SS and Tad systems of the reference dataset (Table S1). In the validation dataset we missed some components of four of the 17 T2SS (Table S2). We could retrieve three of them by modifying the default T2SS model, *e.g.,* the “Xps-type” system could be detected by decreasing the required number components ^53^. Additionally, we missed the very atypical T2SS of *Legionella pneumophila* because it failed the co-localisation criterion (unusually, it is encoded in 5 distant loci, Table S2) ^54^. A default model with more relaxed parameters in terms of co-localisation and sequence similarity would have identified all T2SS, but at the cost of less correct discrimination from the two other systems.

The quality of the default T2SS model was confirmed by the analysis of genomic data. Proteins matched by the protein profiles of T2SS were typically either highly or poorly clustered (Fig. S1a). These two extremes were respectively found to correspond to components of the T2SS and of other systems. The T2SS components co-localised much closer than the imposed distance threshold (d≤5, Fig. S1b). The vast majority (99%) of the T4P were encoded in multiple distant loci, which is accepted but not required by the model whereas most T2SS were encoded in one single locus (96.5%). To verify that the T2SS, T4P, and Tad loci were correctly classed, we compared the HMMER scores of proteins matched by protein profiles from different systems. Proteins matching profiles from two types of systems scored systematically higher for the system in which they were classed even though this is not a classification criterion in the model (Fig. 2b-c). This suggests that our classification is accurate.

We detected 400 T2SS in 360 genomes, 379 T4P in 377 genomes, and 425 Tad pili in 323 genomes. The high abundance of Tad pili is surprising given that they are much less studied than the other systems. Interestingly, we found one Tad pilus with the outer membrane channel (the secretin) in one of the rare Firmicutes with an outer membrane (Clostridia, *Acetohalobium arabaticum* DSM 5501) ^55^, and also in Acidobacteria, Chlorobi, and Nitrospirae. T4P, T2SS, and to a lesser extent Tad pili, were usually found in a single copy per genome, but some genomes encoded up to three systems (Fig. 2d). The observed small number of T2SS per genome reinforces previous suggestions that many T2SS might secrete several different proteins ^56^.

### T3SS and T4SS

T3SS and T4SS are capable of secreting proteins directly into other cells. The T3SS, sometimes also termed non-flagellar T3SS or NF-T3SS, evolved from the flagellar secretion system and is encoded by 15 to 25 genes usually in a single locus ^24,57,58^ (Fig. 3a). Many of the core components of this system are homologous to the distinct F-T3SS that is part of the bacterial flagellum ^59-61^. We have previously proposed models that accurately discriminate between the T3SS and the flagellum ^24^. We used the same models in this work. We identified 434 NF-T3SS in 334 genomes and 837 flagella in 762 genomes. Some genomes encode many T3SS, *e.g., Burkholderia thailandensis* MSMB121 encodes 4 T3SS. These results match experimental data showing that in *Burkholderia pseudomallei* the multiple T3SS target different types of cells ^62^, and that in *Salmonella enterica* the two T3SS are expressed at different moments in the infection cycle (reviewed in ^63^). Multiplicity of T3SS is therefore likely to be associated with complex lifestyles.

T4SS are involved in protein secretion, in conjugation and in some cases in DNA release to, or uptake from, the environment ^64^. Here, we distinguished the protein secretion T4SS from the conjugation-related T4SS, which requires a relaxase ^65^, by naming them respectively pT4SS and cT4SS. It should be noted that some cT4SS are also able to secrete proteins ^64^. We have previously built and validated profiles and models for the pT4SS, and cT4SS ^66^ (Fig. 3). The latter can be divided in eight sub-types corresponding to different mating pair formation complexes (MPF) ^30^, of which six are found in diderm-LPS bacteria, and only two are known to include pT4SS (MPF_I_ and MPF_T_). To test the specificity of the models of each T4SS sub-type, we studied the close co-occurrence of T4SS components. The results show that most protein profiles are highly specific to each T4SS sub-type (Fig. S2). Hence, our profiles are able to identify and distinguish between these different systems. We identified 156 pT4SS (among 990 T4SS) in 130 genomes of diderm bacteria (Table S3).

### T5SS

T5SS are divided in five types (reviewed in ^67-70^). Four types encode the translocator (pore-forming) and the passenger (secreted) domains in a single gene: the classical autotransporter (T5aSS), the trimeric autotransporter (T5cSS), the inverted autotransporter (T5eSS), and the fused two-partner system (T5dSS). In two-partner systems (T5bSS), the translocator and passenger are encoded in two separate (typically contiguous) genes. T5SS rely on the Sec machinery for inner-membrane translocation and require other cellular functions for biogenesis. Many of these functions are ubiquitous in diderm-LPS bacteria and do not facilitate the identification of T5SS. Hence, our models only included information on the conserved, mandatory translocator domain of T5SS (Fig. 4, Fig. S3). Two recently proposed families of T5SS - T5dSS and T5eSS ^71,72^ - were not matched by the T5SS profiles. We will build specific profiles for the detection of these sub-types when enough experimentally validated examples become available.

Our models were able to identify all T5SS in the *reference* and *validation* datasets, with the exception of an atypical previously described T5bSS of *Pseudomonas aeruginosa* consisting of a translocator domain fused with a component of the chaperone usher pathway ^73^. We found 3,829 T5aSS in the genomes of diderm bacteria, which makes them by far the most abundant secretion system in our dataset. Certain *Chlamydiae* genomes contain up to 21 T5aSS. We found 1,125 T5bSS (0-8 per genome) and 849 T5cSS (0-24 per genome). T5SS were encoded in 62% of the genomes of diderm bacteria.

### T6SS

T6SS secrete effectors to bacterial or eukaryotic cells. They were recently divided in three sub-types ^40^, among which T6SS^i^ is by far the most studied ^74-80^. This sub-type has more than a dozen components^77,81^. We built profiles for 14 conserved protein families (Fig. 5a, Materials and Methods, Tables S1, S4, S5), of which 13 were previously described as the most conserved components of the T6SS^i^ ^20^. The remaining profile corresponds to the PAAR-repeat-containing EvpJ protein family of the spike complex^82^, present in eight out of the nine T6SS^i^ in the reference dataset. Using this model we identified all T6SS^i^ of the reference and of the validation datasets. This suggests that the T6SS^i^ model is very accurate. We only found an inaccuracy in *Escherichia coli* O42 where two systems are adjacent in the genome and were identified as a single system. Part of the T6SS^i^ machinery is structurally homologous to the puncturing device of phages, from which it may have originated ^83^. Yet, our model did not identify a T6SS^i^ in any of the 998 phages present in GenBank, showing that it does not mistake puncturing devices for components of the T6SS.

We identified 652 T6SS^i^ in 409 bacterial genomes, with up to 6 T6SS^i^ per genome in some *Burkholderia pseudomallei* strains. Around 9% of the T6SS^i^ are encoded in multiple loci in the genome. Interestingly, 35% of the replicons encoding a T6SS^i^ encode TssI (VgrG) away from the main loci, with either a PAAR-containing component (EvpJ) or the chaperone TssD (Hcp), or both (Fig. 5b-d). PAAR-motifs promote the physical interaction between VgrG and toxins, which are often encoded in the same locus ^80,82,84^. It has recently been proposed that VgrG might also be involved in toxin export in a T6SS-independent way ^80^. Genomes lacking T6SS^i^ do carry some of these small *tssl*-associated clusters, although this corresponds to only 8% of the clusters we could identify. Hence, the identification of loci encoding TssI might uncover new T6SS^i^ effectors.

The T6SS^ii^ sub-type described in *Francisella tularensis,* is involved in subversion of the immune system (growth in macrophages) and virulence ^39,85-87^. Three of the components of the T6SS^ii^ were seldom matched by T6SS^i^ profiles (*tssBCL*), complicating the detection of T6SS^ii^ with the T6SS^i^ model. We built 17 protein profiles and made a specific model for T6SS^ii^ based on a *Francisella tularensis* system (see Fig. 5, Materials and Methods and Table S5) ^87,88^. Using HHsearch ^89^, we confirmed distant homology between the proteins encoded by *tssBCIL* and T6SS^i^ and/or T6SS^iii^ components (p-value<0.001). The model detected 30 T6SS^ii^ in bacterial genomes. All instances were identified exclusively within the 18 genomes of *Francisella,* and all genomes on the genus contained at least one system.

A recent report identified T6SS^iii^ in *Flavobacterium johnsoniae* and *Bacteroides fragilis* and showed it to be involved in bacterial competition ^40^. This sub-type included 9 homologs of the 13 described core components of T6SS^i^ and lacked homologs of the “trans-envelope subcomplex” (Fig. 5). Furthermore, it had three specific components (TssN, TssO and TssP). We built 13 protein profiles based on the analysis of the reported three loci, including the nine homologs to core components of T6SS^i^, the nearly core T6SS^i^ component EvpJ, and two specific components TssN and TssP. We could not build a protein profile for TssO because of the lack of representative sequences similar enough to constitute a protein family (Table S5). The parameters for the models were inferred from the analysis of co-localised hits of the T6SS^iii^ components protein profiles (Fig. S4). We identified 20 T6SS^iii^ in 18 of the 97 Bacteroidetes genomes. TssQ was not previously recognized as conserved but was found in 50% of the systems identified in genomes. It had no PFAM hit, but InterProScan predicted the presence of one secretion signal and cell localisation at the outer-membrane ^90,91^. Interestingly, we could find occurrences of EvpJ (harbouring a PAAR domain) within 6 of the 20 T6SS^iii^ main loci, and outside of the main locus in 6 genomes with a T6SS^iii^. This component co-localised with TssD (Hcp), TssE, or TssD and TssI (VgrG). This suggests it might have similar roles in both T6SS^i^ and T6SS^iii^. T6SS^iii^ was only identified among Bacteroidetes.

### T9SS

A novel protein secretion system, T9SS or PorSS, has been described in *F. johnsoniae* and *Porphyromonas gingivalis* ^9,92^. It is required for the secretion of components of the gliding motility apparatus, adhesins and various hydrolytic enzymes. We used eight protein profiles from TIGRFAM and PFAM for five components (some having several profiles), and built protein profiles for five other components (Tables S4-S5, Materials and Methods). One of the profiles was not specifically associated with T9SS; it is part of the gliding motility machinery (GldJ). It was included in the model because it clusters with some of the T9SS components and thus facilitates their identification. Hence, the model includes 13 protein profiles for 10 core T9SS components ^92^ (Fig. 6, Fig. S5). Four components of the T9SS are scattered in the chromosome, whereas the others are encoded in two gene clusters. We detected 60 T9SS in 60 of the 97 genomes of Bacteroidetes, and none in other clades, as previously shown ^10^. T9SS were found in 62% of the species of Bacteroidetes.

### Distribution of secretion systems

To the best of our knowledge, this is the first report comparing the frequency of all well-known protein secretion systems of diderm-LPS bacteria in bacterial genomes. Therefore, we analysed the distribution of these protein secretion systems in relation to bacterial phylogeny, including clades with more than four genomes and with reliable information on their phylogenetic position (Fig. 7). Only three clades, Alpha-, Beta- and Gamma-Proteobacteria, encoded all the six most-studied protein secretion systems (T1-T6SS^i^). Delta- and Epsilon-Proteobacteria showed fewer or no T2SS, T3SS and pT4SS. Most other clades encode fewer types of systems. The distributions of T3SS, T4SS, T6SS, and T9SS have been described recently ^10,20,24,30,40^, so we shall focus our analysis on the other systems and on their relative distribution.

**Figure 7.**
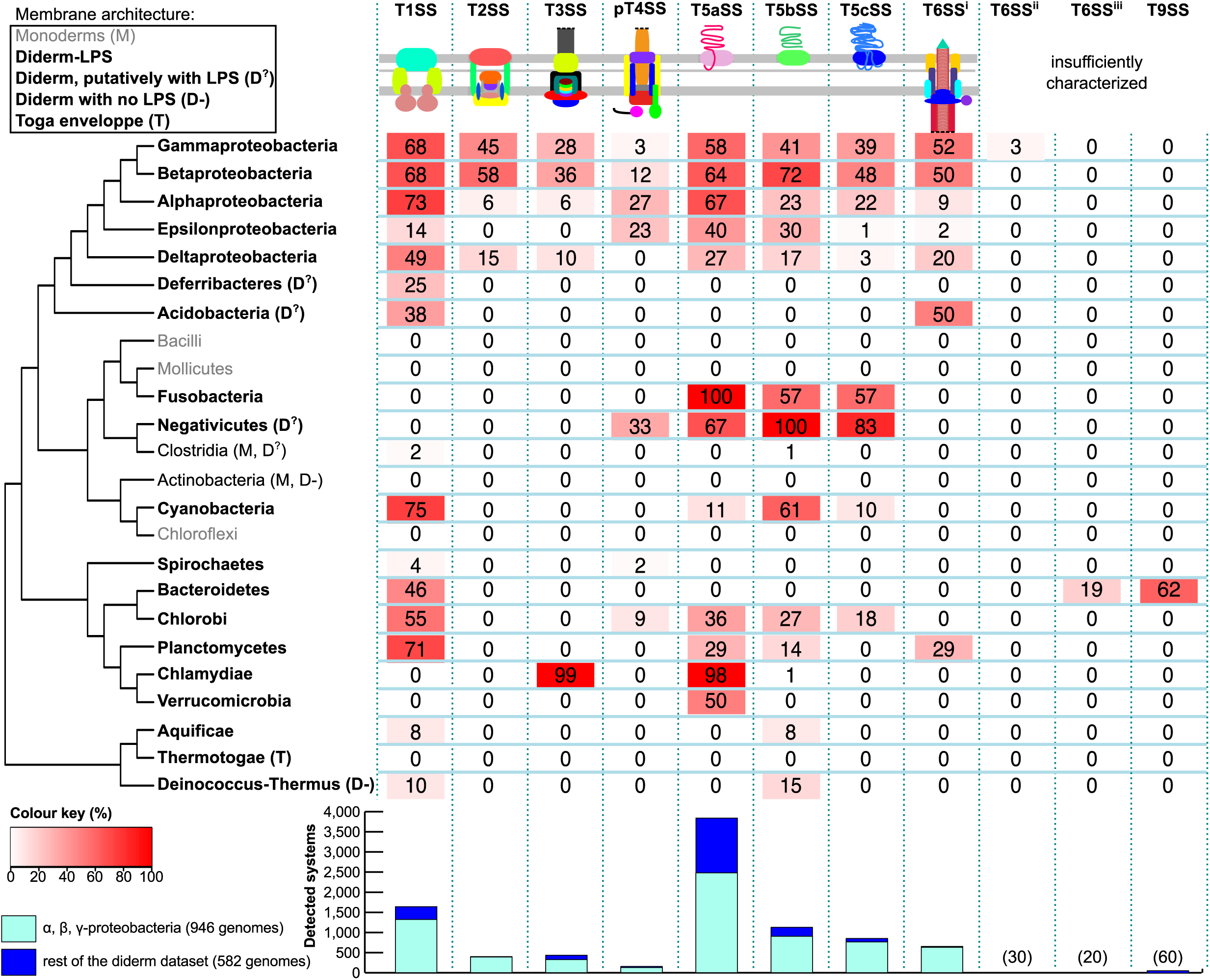
Phylogenetic distribution of protein secretion systems in bacteria. Within each clade, the proportion of genomes harbouring each system is indicated in boxes whose colours follow a gradient from full red (100%) to white (0%) (see legend). Clades were classed as monoderms (grey or “M” symbol), diderms with LPS-containing outer membranes (Diderm-LPS in bold, no symbol), diderms with homologs of LPS pathway that putatively have LPS (D^?^) and diderms with no LPS (D-). The peculiar envelope of the Thermotogae is indicated (T). The Firmicutes are typically monoderms, but some of their members are diderms (the Negativicutes, some Clostridia, Mycobacteria). The bar plot shows the number of detected systems. Bars are split in two categories to separate on one side Alpha- Beta- and Gamma-proteobacteria, and on the other genomes from other bacteria. We display the number of occurrences of systems occurring rarely in our dataset on top of the bars. Clades with less than 4 genomes and/or with unreported phylogenetic position are not shown (*i.e.,* Chrysiogenetes, Gemmatimonadetes, Nitrospirae and Thermodesulfobacteria). This sketch tree was drawn from the compilation of different published phylogenetic analyses ^114-117^

T1SS and T5SS are the most widespread protein secretion systems (Fig. 7, see below). We identified T1SS in phyla as diverse as Spirochaetes, Planctomycetes, Aquificae, Bacteroidetes, and Cyanobacteria. T1SS involved in the secretion of glycolipids for heterocysts formation were recently described in filamentous Cyanobacteria ^93,94^. We find that T1SS are particularly abundant in this clade, where we identified 156 systems in 72 genomes, with 75% of the genomes harbouring at least one T1SS. The three types of T5SS showed similar taxonomic distributions, even if T5cSS are less widespread (Fig. 7). Some phyla include only one type of T5SS: T5aSS in Thermodesulfobacteria and T5bSS in Aquificae and Deinococcus-Thermus. These clades have too few representative genomes to conclude if they lack the other T5SS.

We identified very few T2SS outside Proteobacteria. T2SS were also absent from the 98 genomes of Epsilon-proteobacteria. We did find a T2SS in a non-Proteobacteria, *Desulfurispirillum indicum* S5, a free-living spiral-shaped aquatic Chrysiogenetes (also encoding a T1SS). We could not find a description of the membrane architecture for this species, but our analysis reinforces previous suggestions that it is a diderm ^95^. Putative T2SS were previously identified in clades where we failed to identify complete systems: *Synechococcus elongatus* (Cyanobacteria), *Chlamydia trachomatis* (Chlamydiae) and *Leptospira interrogans* (Spirochaetes) ^28,96-98^. The cyanobacterial system, which has a role in protein secretion and biofilm formation, seems to be a typical T4P encoded in multiple loci. The role of T4P in secreting proteins that are not part of its structure has been described before ^99^. To the best of our knowledge, the function of the *Leptospira* system was not experimentally tested. The Chlamydiae system was indeed associated with protein secretion ^97^. From the point of view of our models the putative T2SS from these two last clades form incomplete systems (although they could be retrieved by lowering the minimum required number of components for a valid T2SS in the model). Preliminary phylogenetic analyses did not allow conclusive assignment of these systems to T2SS or to T4P. Further experimental and computational work will be necessary for their precise characterisation.

### Secretion systems and the cell envelope

The distribution of secretion systems is linked with the cell envelope. Expectedly, all genomes of monoderms lacked loci encoding diderm-like protein secretion system. Several clades of diderm bacteria have few types of protein secretion systems, but only one lacks them all: the Thermotogae. These bacteria are thermophilic, and one could speculate that high temperatures could be incompatible with protein secretion systems of mesophiles. Yet, life under high temperatures is also typical of the sister-clade Aquificae, which has T1SS and T5SS. Instead, the lack of typical protein secretion systems in Thermotogae might be caused by the peculiar sheath-like structure present in their outer cell envelope, the “toga” ^100^. This may have led to the evolution of different secretion systems. Accordingly, only a few porins have been identified so far in Thermotogae ^101^. In an analogous way, *Mycobacteria* (Actinobacteria), which have a peculiar mycolate outer membrane, have specific secretion systems ^12^.

The cell envelope architecture is a key determinant of the evolution of secretion systems in other clades as well. The extracellular structures of T3SS are tightly linked with the type of eukaryote cell (plant vs. animal) with which the bacterium interacts ^24^. Sub-types of T4SS are also specific to certain bacterial cell envelopes ^30^. Interestingly, some diderm bacteria from taxa dominated by monoderms have protein secretion systems homologous to those of Proteobacteria (including Clostridia, Cyanobacteria, Fusobacteria and Negativicutes). For example, we identified in Negativicutes (a clade of Firmicutes) putative pT4SS and the three types of T5SS. Some genomes of Halanaerobiales (a sub-clade of Clostridia, Firmicutes) encode T1SS and T5bSS. Similarities in the cell envelope may thus lead to the presence of similar systems in very distant bacteria.

### Conclusion

Using our models, we were able to identify nearly all protein secretion systems in both the *reference* and the *validation* datasets. The few systems we have missed are very atypical (such as the scattered T2SS of *Legionella*) or have homologous components in other types of systems (several T2SS). In the latter case, the relaxation of the parameters of the T2SS model allowed their identification. We emphasize that our models are publicly available and can be modified by the user to increase sensitivity, for example by simply relaxing the parameters for components detection (HMMER i-evalue and profile coverage), or by altering the parameters of the model *e.g.,* by decreasing the minimal number of components required for a valid system or by relaxing the co-localisation criterion. Nevertheless, this will inevitably lead to an increased number of false positives. Complementary analyses can also facilitate the identification of systems. For example, the results of Fig. 2b-c show that the comparison of the scores of the hits of the protein profiles discriminate between homologous components in different systems, even when the information on genetic organization is lacking or insufficient. This is one of the advantages of using specifically designed protein profiles, instead of generic profiles as can be found in PFAM: the system-specific profiles distinguish between homologs components in different types of molecular systems. Comparisons of the scores for homologous proteins, or phylogenetic analyses could also be useful to analyse poorly assembled genomic or metagenomic data where co-localisation data is unavailable.

Rapid evolution of extracellular components of protein secretion systems and the paucity of experimentally studied systems outside Proteobacteria may have led us to under-estimate their presence in less studied clades. Yet, several pieces of evidence suggest that we may have not missed many homologous systems. 1) In most cases we identified all known systems in our *reference* and *validation* datasets. 2) We identified at least one type of secretion system in almost all clades of diderm bacteria. 3) We identified components of T4P and Tad (homologous to T2SS), F-T3SS (homologous to NF-T3SS), and cT4SS (homologous to pT4SS) with certain profiles for the protein secretion systems in many clades, including monoderms (Table S3). If one is able to pinpoint systems that are so distantly related, one should have been able to identify most systems of a given type. Instead, it is tempting to speculate that in clades other than Proteobacteria there are alternative and currently unknown protein secretion systems. Interestingly, the recently discovered T6SS^iii^ and T9SS are restricted to Bacteroidetes in our dataset ^9,40^, while T6SS^ii^ are only found in *Francisella.* The absence of well-known protein secretion systems in certain taxa and the presence of lineage-specific systems are consistent with the hypothesis that novel systems remain to be discovered.

## Materials and Methods

### Data

The genomes of bacteria (2,484) and archaea (159) were downloaded from NCBI RefSeq (ftp://ftp.ncbi.nih.gov/genomes/, November 2013). From these, we sorted out 1,528 genomes of bacteria classed as diderm in the literature ^102-104^ (Table S3). A total of 998 genomes of phages were downloaded from Genbank (last access, February 2013). The sequences of the reference protein secretion systems were retrieved from Genbank or from complete genomes (Tables S1 and S2).

### Systems definition and identification

We built a dataset of experimentally studied secretion systems (T1SS-T6SS, T9SS) and related assemblages (Tad, T4P and the bacterial flagellum) from the analysis of published data. This reference dataset was used to build the models of each type of system (Table S1) using MacSyFinder. This software is publicly available and was described in detail previously ^25^. Here, we focus on the features of the program that are pertinent for the secretion systems models. A model in MacSyFinder defines the components of the secretion system, the minimal acceptable number of components and their genetic organisation. Among other things (see http://macsyfinder.readthedocs.org for full documentation), one can specify the following relevant information. 1) Systems are encoded in a single locus (*single-locus* system) or in several loci (*scattered* system). 2) Core components (ubiquitous and essential) are defined as *mandatory.* 3) Components that are accessory or poorly conserved in sequence are defined as *accessory*. These components are accessory for the computational model, but their function may be essential. This happens when different proteins have analogous functions or when proteins evolve so fast that distant homologs are not recognisable. 4) Some genes are ubiquitous and specific to a system and can be defined as forbidden in models of other systems. This facilitates discrimination between systems with homologous components. For example, the NF-T3SS-specific secretin may be declared as forbidden in the F-T3SS. 5) An occurrence of a system is validated when a pre-defined number (*quorum*) of mandatory components and/or sum of mandatory and accessory components is found ^25^. 6) Components can be defined as reciprocally *exchangeable* in the quorum (which prevents them from being counted twice). 7) Two components are co-localised when they are separated by less than a given number of genes (parameter *d=inter_gene_max_space*). 8) A component defined with the *loner* attribute does not need to be co-localised with other components to be part of a system (*e.g.,* OMF in T1SS). 9) A component that can participate in several instances of a system (*e.g.,* OMF in T1SS) receives the *multi_system* attribute. These different properties can be combined when necessary.

The models for the different protein secretion systems were described in files following a dedicated XML grammar ^25^ and named after the system (*e.g.,* T1SS.xml, File S1). Models can be easily modified on the standalone version of MacSyFinder. The webserver allows the modification of the most important search parameters.

MacSyFinder was used to identify protein secretion systems in bacterial genomes in several steps (for a full description of the software see ^25^). The program did the search in three steps. Firstly, components were identified using protein profile searches with HMMER ^41^. The hits were kept when the alignments covered more than 50% of the protein profile and the i-evalue<10^−3^ (default parameters). Secondly, the components were clustered according to their proximity in the genome using the parameter d. Finally, the clusters were validated if they passed the criteria in the models of the systems.

### Definition of protein profiles

The models included 204 protein profiles. Among these, the two profiles for T5aSS and T5cSS were extracted from PFAM ^67,90^, and eight profiles for T9SS were extracted from PFAM or TIGRFAM ^90,105^. The remaining 194 were the result of our previous work ^24,66,106^ or this study (84 protein profiles for T1SS, T2SS, Tad, type IV pilus, T5bSS, T6SS^i^, T6SS^ii^, T6SS^iii^ and T9SS, listed in Table S4). To build these profiles we sampled the experimentally studied systems for proteins representative of each component of each system. We selected examples within these to maximise sequence diversity (Table S1). Protein families were constructed by clustering homologous proteins. The details of the methods and parameters used to build each protein profile are described in Table S5. In the case of the T9SS, where only two systems were experimentally characterised, we used components from the well-studied system of *F. johnsoniae* (or *P. gingivalis* when the gene was absent from *F. jonhsoniae*) for Blastp searches against our database of complete genomes, and retained the best sequence hits (e-value < 10^−20^) to constitute protein families. A similar approach was taken to build protein profiles for the T6SS^ii^, based on the *Francisella tularensis subsp. tularensis* SCHU S4 FPI system displayed in Table 1 of ^39^. The largest families obtained were aligned and manually curated to produce hidden Markov model profiles with HMMER 3.0 ^41^.

### Availability

Detection and visualization of all systems described in this paper can be performed online on the Mobyle-based ^107^ webserver TXSScan: http://mobyle.pasteur.fr/cgi-bin/portal.py#forms::txsscan. Detection can also be performed locally using the standalone program MacSyFinder ^25^, and the sets of models and profiles described here. MacSyFinder is freely available for all platforms at https://github.com/gem-pasteur/macsyfinder. Models and required protein profiles are available as supplemental material (File S1) at https://research.pasteur.fr/en/tool/txsscan-models-and-profiles-for-protein-secretion-systems (https://researchpullzone-yhello.netdna-ssl.com/wp-content/uploads/2015/10/research.pasteur.fr_txsscan-models-and-profiles-for-protein-secretion-systems.gz) The models are provided as simple text (XML) files, so they can be easily modified and extended by the user. The results of MacSyFinder can be visualized with MacSyView, that is available online at http://macsyview.web.pasteur.fr or for download at https://github.com/gem-pasteur/macsyview (also included in the release of MacSyFinder).

## Acknowledgments

We are grateful to Elie Dassa, Olivera Francetic, and Marc Garcia-Garcerà for fruitful discussions, and Hervé Ménager for its contribution to MacSyView. We thank Eric Duchaud for discussions and comments on T9SS. This work was supported by the CNRS, the Institut Pasteur and the European Research Council (grant EVOMOBILOME, number 281605). JC is a member of the French doctoral school “Ecole Doctorale Interdisciplinaire Européenne Frontières du Vivant ED474”.

## Authors’ contributions

SSA and EPCR designed the analyses. SSA designed the secretion systems models and profiles, and performed the analyses. JC and JG contributed to the T4SS models, the T4SS protein profiles and the corresponding analyses. BN contributed to the MacSyFinder and MacSyView software, and created the online interface for TXSScan and MacSyView. MT contributed to the T9SS models and the T9SS protein profiles. SSA and EPCR wrote the manuscript with the help of the other authors. All authors read and approved the full manuscript.

**Abbreviations**
ESX: ESAT-6 secretion system
HMM: hidden Markov model
LPS: Lipopolysaccharide
OMF: outer-membrane porin of T1SS
MFP: membrane-fusion protein from T1SS.
MFS: major facilitator superfamily
MPF: mating pair formation complex
RND: resistance nodulation cell division
T1SS: Type I secretion system
T2SS: Type II secretion system
T3SS: Type III secretion system
T4P: Type IV pilus
T4SS: Type IV secretion system
cT4SS: conjugative T4SS
pT4SS: protein delivery T4SS
T5SS: Type V secretion system
T6SS: Type VI secretion system
T9SS: Type IX secretion system
Tad: Tight adherence
XML: Extensible Markup Language

## References

1. Wandersman C & Delepelaire P. Bacterial iron sources: from siderophores to hemophores. Annu. Rev. Microbiol. 58, 611 (2004).

2. Ruhe ZC, Low DA & Hayes CS. Bacterial contact-dependent growth inhibition. Trends Microbiol. 21, 230 (2013).

3. Viprey V, Del Greco A, Golinowski W, Broughton WJ & Perret X. Symbiotic implications of type III protein secretion machinery in Rhizobium. Mol. Microbiol. 28, 1381 (1998).

4. Ma W & Guttman DS. Evolution of prokaryotic and eukaryotic virulence effectors. Curr. Opin. Plant Biol. 11, 412 (2008).

5. Raymond B et al. Subversion of trafficking, apoptosis, and innate immunity by type III secretion system effectors. Trends Microbiol. 21, 430 (2013).

6. Bleves S et al. Protein secretion systems in Pseudomonas aeruginosa: A wealth of pathogenic weapons. Int. J. Med. Microbiol. 300, 534 (2010).

7. Dalbey RE & Kuhn A. Protein traffic in Gram-negative bacteria--how exported and secreted proteins find their way. FEMS Microbiol. Rev. 36, 1023 (2012).

8. Chang JH, Desveaux D & Creason AL. The ABCs and 123s of bacterial secretion systems in plant pathogenesis. Annu Rev Phytopathol 52, 317 (2014).

9. Sato K et al. A protein secretion system linked to bacteroidete gliding motility and pathogenesis. Proc. Natl. Acad. Sci. U. S. A. 107, 276 (2010).

10. McBride MJ & Zhu Y. Gliding motility and Por secretion system genes are widespread among members of the phylum bacteroidetes. J. Bacteriol. 195, 270 (2013).

11. Desvaux M, Hebraud M, Talon R & Henderson IR. Secretion and subcellular localizations of bacterial proteins: a semantic awareness issue. Trends Microbiol. 17, 139 (2009).

12. Abdallah AM et al. Type VII secretion-mycobacteria show the way. Nat. Rev. Microbiol. 5, 883 (2007).

13. Tseng TT, Tyler BM & Setubal JC. Protein secretion systems in bacterial-host associations, and their description in the Gene Ontology. BMC Microbiol. 9 Suppl 1, S2 (2009).

14. Planet PJ, Kachlany SC, DeSalle R & Figurski DH. Phylogeny of genes for secretion NTPases: identification of the widespread tadA subfamily and development of a diagnostic key for gene classification. Proc. Natl. Acad. Sci. U. S. A. 98, 2503 (2001).

15. Minamino T & Namba K. Self-assembly and type III protein export of the bacterial flagellum. J. Mol. Microbiol. Biotechnol. 7, 5 (2004).

16. Pelicic V. Type IV pili: e pluribus unum? Mol. Microbiol. 68, 827 (2008).

17. Peabody CR et al. Type II protein secretion and its relationship to bacterial type IV pili and archaeal flagella. Microbiology 149, 3051 (2003).

18. Nogueira T, Touchon M & Rocha EP. Rapid evolution of the sequences and gene repertoires of secreted proteins in bacteria. PLoS ONE 7, e49403 (2012).

19. Gophna U, Ron EZ & Graur D. Bacterial type III secretion systems are ancient and evolved by multiple horizontal-transfer events. Gene 312, 151 (2003).

20. Boyer F, Fichant G, Berthod J, Vandenbrouck Y & Attree I. Dissecting the bacterial type VI secretion system by a genome wide in silico analysis: what can be learned from available microbial genomic resources? BMC Genomics 10, 104 (2009).

21. Ren CP et al. The ETT2 gene cluster, encoding a second type III secretion system from Escherichia coli, is present in the majority of strains but has undergone widespread mutational attrition. J. Bacteriol. 186, 3547 (2004).

22. Huynen M, Snel B, Lathe W, 3rd & Bork P. Predicting protein function by genomic context: quantitative evaluation and qualitative inferences. Genome Res. 10, 1204 (2000).

23. Wolf YI, Rogozin IB, Kondrashov AS & Koonin EV. Genome alignment, evolution of prokaryotic genome organization, and prediction of gene function using genomic context. Genome Res. 11, 356 (2001).

24. Abby SS & Rocha EP. The non-flagellar type III secretion system evolved from the bacterial flagellum and diversified into host-cell adapted systems. PLoS Genet. 8, e1002983 (2012).

25. Abby SS, Neron B, Menager H, Touchon M & Rocha EP. MacSyFinder: A Program to Mine Genomes for Molecular Systems with an Application to CRISPR-Cas Systems. PLoS ONE 9, e110726 (2014).

26. Yen MR et al. Protein-translocating outer membrane porins of Gram-negative bacteria. Biochim. Biophys. Acta 1562, 6 (2002).

27. Pallen MJ, Beatson SA & Bailey CM. Bioinformatics, genomics and evolution of non-flagellar type-III secretion systems: a Darwinian perspective. FEMS Microbiol. Rev. 29, 201 (2005).

28. Cianciotto NP. Type II secretion: a protein secretion system for all seasons. Trends Microbiol. 13, 581 (2005).

29. Saier MH, Ma CH, Rodgers L, Tamang DG & Yen MR. Protein secretion and membrane insertion systems in bacteria and eukaryotic organelles. Adv. Appl. Microbiol. 65, 141 (2008).

30. Guglielmini J, de la Cruz F & Rocha EP. Evolution of conjugation and type IV secretion systems. Mol. Biol. Evol. 30, 315 (2013).

31. Barret M, Egan F & O’Gara F. Distribution and diversity of bacterial secretion systems across metagenomic datasets. Environmental microbiology reports 5, 117 (2013).

32. Delepelaire P. Type I secretion in gram-negative bacteria. Biochim. Biophys. Acta 1694, 149 (2004).

33. Bouige P, Laurent D, Piloyan L & Dassa E. Phylogenetic and functional classification of ATP-binding cassette (ABC) systems. Curr. Protein Pept. Sci. 3, 541 (2002).

34. Dassa E & Bouige P. The ABC of ABCS: a phylogenetic and functional classification of ABC systems in living organisms. Res. Microbiol. 152, 211 (2001).

35. Jacob-Dubuisson F, Fernandez R & Coutte L. Protein secretion through autotransporter and two-partner pathways. Biochim. Biophys. Acta 1694, 235 (2004).

36. Henderson IR, Navarro-Garcia F, Desvaux M, Fernandez RC & Ala’Aldeen D. Type V protein secretion pathway: the autotransporter story. Microbiol. Mol. Biol. Rev. 68, 692 (2004).

37. Linke D, Riess T, Autenrieth IB, Lupas A & Kempf VA. Trimeric autotransporter adhesins: variable structure, common function. Trends Microbiol. 14, 264 (2006).

38. Tomich M, Planet PJ & Figurski DH. The tad locus: postcards from the widespread colonization island. Nat. Rev. Microbiol. 5, 363 (2007).

39. Broms JE, Sjostedt A & Lavander M. The Role of the Francisella Tularensis Pathogenicity Island in Type VI Secretion, Intracellular Survival, and Modulation of Host Cell Signaling. Front. Microbiol. 1, 136 (2010).

40. Russell AB et al. A Type VI Secretion-Related Pathway in Bacteroidetes Mediates Interbacterial Antagonism. Cell Host Microbe 16, 227 (2014).

41. Eddy SR. Accelerated Profile HMM Searches. PLoS Comput. Biol. 7, e1002195 (2011).

42. Eddy SR. Profile hidden Markov models. Bioinformatics 14, 755 (1998).

43. Holland IB, Schmitt L & Young J. Type 1 protein secretion in bacteria, the ABC-transporter dependent pathway. Mol. Membr. Biol. 22, 29 (2005).

44. Kanonenberg K, Schwarz CK & Schmitt L. Type I secretion systems - a story of appendices. Res. Microbiol. 164, 596 (2013).

45. Paulsen IT, Park JH, Choi PS & Saier MH, Jr. A family of gram-negative bacterial outer membrane factors that function in the export of proteins, carbohydrates, drugs and heavy metals from gram-negative bacteria. FEMS Microbiol. Lett. 156, 1 (1997).

46. Dinh T, Paulsen IT & Saier MH, Jr. A family of extracytoplasmic proteins that allow transport of large molecules across the outer membranes of gram-negative bacteria. J. Bacteriol. 176, 3825 (1994).

47. Dassa E. Natural history of ABC systems: not only transporters. Essays Biochem. 50, 19 (2011).

48. Davidson AL, Dassa E, Orelle C & Chen J. Structure, function, and evolution of bacterial ATP-binding cassette systems. Microbiol. Mol. Biol. Rev. 72, 317 (2008).

49. Koronakis V, Eswaran J & Hughes C. Structure and function of TolC: the bacterial exit duct for proteins and drugs. Annu. Rev. Biochem. 73, 467 (2004).

50. Burland V et al. The complete DNA sequence and analysis of the large virulence plasmid of Escherichia coli O157:H7. Nucleic Acids Res. 26, 4196 (1998).

51. Korotkov KV, Sandkvist M & Hol WG. The type II secretion system: biogenesis, molecular architecture and mechanism. Nat. Rev. Microbiol. 10, 336 (2012).

52. Nivaskumar M & Francetic O. Type II secretion system: a magic beanstalk or a protein escalator. Biochim. Biophys. Acta 1843, 1568 (2014).

53. Karaba SM, White RC & Cianciotto NP. Stenotrophomonas maltophilia Encodes a Type II Protein Secretion System That Promotes Detrimental Effects on Lung Epithelial Cells. Infect. Immun. 81, 3210 (2013).

54. Cianciotto NP. Many substrates and functions of type II secretion: lessons learned from Legionella pneumophila. Future Microbiol. 4, 797 (2009).

55. Zhilina TN & Zavarzin GA. Extremely halophilic, methylotrophic, anaerobic bacteria. FEMS Microbiol. Lett. 87, 315 (1990).

56. Rondelet A & Condemine G. Type II secretion: the substrates that won’t go away. Res. Microbiol. 164, 556 (2013).

57. Galan JE & Wolf-Watz H. Protein delivery into eukaryotic cells by type III secretion machines. Nature 444, 567 (2006).

58. Cornelis GR. The type III secretion injectisome. Nat. Rev. Microbiol. 4, 811 (2006).

59. Ginocchio CC, Olmsted SB, Wells CL & Galan JE. Contact with epithelial cells induces the formation of surface appendages on Salmonella typhimurium. Cell 76, 717 (1994).

60. Van Gijsegem F et al. The hrp gene locus of Pseudomonas solanacearum, which controls the production of a type III secretion system, encodes eight proteins related to components of the bacterial flagellar biogenesis complex. Mol. Microbiol. 15, 1095 (1995).

61. Young GM, Schmiel DH & Miller VL. A new pathway for the secretion of virulence factors by bacteria: the flagellar export apparatus functions as a protein-secretion system. Proc. Natl. Acad. Sci. U. S. A. 96, 6456 (1999).

62. Sun GW & Gan YH. Unraveling type III secretion systems in the highly versatile Burkholderia pseudomallei. Trends Microbiol. 18, 561 (2010).

63. Hansen-Wester I & Hensel M. Salmonella pathogenicity islands encoding type III secretion systems. Microbes Infect. 3, 549 (2001).

64. Alvarez-Martinez CE & Christie PJ. Biological diversity of prokaryotic type IV secretion systems. Microbiol. Mol. Biol. Rev. 73, 775 (2009).

65. de la Cruz F, Frost LS, Meyer RJ & Zechner E. Conjugative DNA Metabolism in Gram-negative Bacteria. FEMS Microbiol. Rev. 34, 18 (2010).

66. Guglielmini J et al. Key components of the eight classes of type IV secretion systems involved in bacterial conjugation or protein secretion. Nucleic Acids Res. (2014).

67. Dautin N & Bernstein HD. Protein Secretion in Gram-Negative Bacteria via the Autotransporter Pathway. Annu. Rev. Microbiol. 61, 89 (2007).

68. Mazar J & Cotter PA. New insight into the molecular mechanisms of two-partner secretion. Trends Microbiol. 15, 508 (2007).

69. Leyton DL, Rossiter AE & Henderson IR. From self sufficiency to dependence: mechanisms and factors important for autotransporter biogenesis. Nat. Rev. Microbiol. 10, 213 (2012).

70. Leo JC, Grin I & Linke D. Type V secretion: mechanism(s) of autotransport through the bacterial outer membrane. Philos. Trans. R. Soc. Lond. B. Biol. Sci. 367, 1088 (2012).

71. Salacha R et al. The Pseudomonas aeruginosa patatin-like protein PlpD is the archetype of a novel Type V secretion system. Environ. Microbiol. 12, 1498 (2010).

72. Oberhettinger P et al. Intimin and invasin export their C-terminus to the bacterial cell surface using an inverse mechanism compared to classical autotransport. PLoS ONE 7, e47069 (2012).

73. Ruer S, Ball G, Filloux A & de Bentzmann S. The ‘P-usher’, a novel protein transporter involved in fimbrial assembly and TpsA secretion. EMBOJ. 27, 2669 (2008).

74. Mougous JD et al. A virulence locus of Pseudomonas aeruginosa encodes a protein secretion apparatus. Science 312, 1526 (2006).

75. Hood RD et al. A type VI secretion system of Pseudomonas aeruginosa targets a toxin to bacteria. Cell Host Microbe 7, 25 (2010).

76. Schwarz S et al. Burkholderia type VI secretion systems have distinct roles in eukaryotic and bacterial cell interactions. PLoS Pathog. 6, e1001068 (2010).

77. Silverman JM, Brunet YR, Cascales E & Mougous JD. Structure and regulation of the type VI secretion system. Annu. Rev. Microbiol. 66, 453 (2012).

78. Basler M, Ho BT & Mekalanos JJ. Tit-for-tat: type VI secretion system counterattack during bacterial cell-cell interactions. Cell 152, 884 (2013).

79. Brunet YR, Espinosa L, Harchouni S, Mignot T & Cascales E. Imaging type VI secretion-mediated bacterial killing. Cell reports 3, 36 (2013).

80. Hachani A, Allsopp LP, Oduko Y & Filloux A. The VgrG proteins are “A la carte “ delivery systems for bacterial type VI effectors. J. Biol. Chem. 289, 17872 (2014).

81. Cascales E & Cambillau C. Structural biology of type VI secretion systems. Philos. Trans. R. Soc. Lond. B. Biol. Sci. 367, 1102 (2012).

82. Shneider MM et al. PAAR-repeat proteins sharpen and diversify the type VI secretion system spike. Nature 500, 350 (2013).

83. Pukatzki S, Ma AT, Revel AT, Sturtevant D & Mekalanos JJ. Type VI secretion system translocates a phage tail spike-like protein into target cells where it cross-links actin. Proc. Natl. Acad. Sci. U. S. A. 104, 15508 (2007).

84. Whitney JC et al. Genetically distinct pathways guide effector export through the type VI secretion system. Mol. Microbiol. 92, 529 (2014).

85. Nano FE et al. A Francisella tularensis pathogenicity island required for intramacrophage growth. J. Bacteriol. 186, 6430 (2004).

86. Ludu JS et al. The Francisella pathogenicity island protein PdpD is required for full virulence and associates with homologues of the type VI secretion system. J. Bacteriol. 190, 4584 (2008).

87. Barker JR et al. The Francisella tularensis pathogenicity island encodes a secretion system that is required for phagosome escape and virulence. Mol. Microbiol. 74, 1459 (2009).

88. Camacho C et al. BLAST+: architecture and applications. BMC Bioinformatics 10, 421 (2009).

89. Soding J. Protein homology detection by HMM-HMM comparison. Bioinformatics 21, 951 (2005).

90. Finn RD et al. The Pfam protein families database. Nucleic Acids Res. 36, D281 (2008).

91. Quevillon E et al. InterProScan: protein domains identifier. Nucleic Acids Res. 33, W116 (2005).

92. Kharade SS & McBride MJ. Flavobacterium johnsoniae PorV is required for secretion of a subset of proteins targeted to the type IX secretion system. J. Bacteriol. 197, 147 (2015).

93. Moslavac S et al. A TolC-like protein is required for heterocyst development in Anabaena sp. strain PCC 7120. J. Bacteriol. 189, 7887 (2007).

94. Staron P, Forchhammer K & Maldener I. Structure-function analysis of the ATP-driven glycolipid efflux pump DevBCA reveals complex organization with TolC/HgdD. FEBS Lett. 588, 395 (2014).

95. Rauschenbach I, Yee N, Haggblom MM & Bini E. Energy metabolism and multiple respiratory pathways revealed by genome sequencing of Desulfurispirillum indicum strain S5. Environ. Microbiol. 13, 1611 (2011).

96. Zeng L et al. Extracellular proteome analysis of Leptospira interrogans serovar Lai. Omics : a journal of integrative biology 17, 527 (2013).

97. Nguyen BD & Valdivia RH. Virulence determinants in the obligate intracellular pathogen Chlamydia trachomatis revealed by forward genetic approaches. Proc. Natl. Acad. Sci. U. S. A. 109, 1263 (2012).

98. Schatz D et al. Self-suppression of biofilm formation in the cyanobacterium Synechococcus elongatus. Environ. Microbiol. 15, 1786 (2013).

99. Hager AJ et al. Type IV pili-mediated secretion modulates Francisella virulence. Mol. Microbiol. 62, 227 (2006).

100. Huber R et al. Thermotoga maritima sp. nov. represents a new genus of unique extremely thermophilic eubacteria growing up to 90 C. Arch. Microbiol. 144, 324 (1986).

101. Petrus AK et al. Genes for the major structural components of Thermotogales species’ togas revealed by proteomic and evolutionary analyses of OmpA and OmpB homologs. PLoS ONE 7, e40236 (2012).

102. Sutcliffe IC. A phylum level perspective on bacterial cell envelope architecture. Trends Microbiol. 18, 464 (2010).

103. Francke C et al. Comparative analyses imply that the enigmatic Sigma factor 54 is a central controller of the bacterial exterior. BMC Genomics 12, 385 (2011).

104. Vesth T et al. Veillonella, Firmicutes: Microbes disguised as Gram negatives. Stand Genomic Sci 9, 431 (2013).

105. Haft DH et al. TIGRFAMs and Genome Properties in 2013. Nucleic Acids Res. 41, D387 (2013).

106. Guglielmini J, Quintais L, Garcillan-Barcia MP, de la Cruz F & Rocha EP. The Repertoire of ICE in Prokaryotes Underscores the Unity, Diversity, and Ubiquity of Conjugation. PLoS Genet. 7, e1002222 (2011).

107. Neron B et al. Mobyle: a new full web bioinformatics framework. Bioinformatics 25, 3005 (2009).

108. Nunn DN & Lory S. Product of the Pseudomonas aeruginosa gene pilD is a prepilin leader peptidase. Proc. Natl. Acad. Sci. U. S. A. 88, 3281 (1991).

109. Pepe CM, Eklund MW & Strom MS. Cloning of an Aeromonas hydrophila type IV pilus biogenesis gene cluster: complementation of pilus assembly functions and characterization of a type IV leader peptidase/N-methyltransferase required for extracellular protein secretion. Mol. Microbiol. 19, 857 (1996).

110. Marsh JW & Taylor RK. Identification of the Vibrio cholerae type 4 prepilin peptidase required for cholera toxin secretion and pilus formation. Mol. Microbiol. 29, 1481 (1998).

111. Christie PJ. Type IV secretion: the Agrobacterium VirB/D4 and related conjugation systems. Biochim. Biophys. Acta 1694, 219 (2004).

112. Nagai H & Kubori T. Type IVB Secretion Systems of Legionella and Other Gram-Negative Bacteria. Front. Microbiol. 2, 136 (2011).

113. Schroder G, Schuelein R, Quebatte M & Dehio C. Conjugative DNA transfer into human cells by the VirB/VirD4 type IV secretion system of the bacterial pathogen Bartonella henselae. Proc. Natl. Acad. Sci. U. S. A. 108, 14643 (2011).

114. Abby SS, Tannier E, Gouy M & Daubin V. Lateral gene transfer as a support for the tree of life. Proc. Natl. Acad. Sci. U. S. A. 109, 4962 (2012).

115. Yutin N, Puigbo P, Koonin EV & Wolf YI. Phylogenomics of prokaryotic ribosomal proteins. PLoS ONE 7, e36972 (2012).

116. Wu D et al. A phylogeny-driven genomic encyclopaedia of Bacteria and Archaea. Nature 462, 1056 (2009).

117. Boussau B, Gueguen L & Gouy M. Accounting for horizontal gene transfers explains conflicting hypotheses regarding the position of aquificales in the phylogeny of Bacteria. BMC Evol. Biol. 8, 272 (2008).

118. Souza RC et al. AtlasT4SS: a curated database for type IV secretion systems. BMC Microbiol. 12, 172 (2012).

119. Bi D et al. SecReT4: a web-based bacterial type IV secretion system resource. Nucleic Acids Res. 41, D660 (2013).

120. Li J et al. SecReT6: a web-based resource for type VI secretion systems found in bacteria. Environ. Microbiol. 17, 2196 (2015).

121. Pundhir S & Kumar A. SSPred: A prediction server based on SVM for the identification and classification of proteins involved in bacterial secretion systems. Bioinformation 6, 380 (2011).

122. Martinez-Garcia PM, Ramos C & Rodriguez-Palenzuela P. T346Hunter: a novel web-based tool for the prediction of type III, type IV and type VI secretion systems in bacterial genomes. PLoS ONE 10, e0119317 (2015).

123. Wang Y, Huang H, Sun M, Zhang Q & Guo D. T3DB: an integrated database for bacterial type III secretion system. BMC Bioinformatics 13, 66 (2012).

